# Are 150 km of open sea enough? Gene flow and population differentiation in a bat-pollinated columnar cactus

**DOI:** 10.1101/2023.02.28.530409

**Authors:** Sebastián Arenas, Alberto Búrquez, Enriquena Bustamante, Enrique Scheinvar, Luis E. Eguiarte

## Abstract

Genetic differentiations and phylogeographical patterns are controlled by the interplay between spatial isolation and gene flow. To test the extent of gene flow across an oceanic barrier, we explored the effect of the separation of the peninsula of Baja California on the evolution of mainland and peninsular populations of the long-lived columnar cactus *Stenocereus thurberi*. We analyzed twelve populations throughout the OPC distribution range to assess genetic diversity and structure using chloroplast DNA sequences. Genetic diversity was higher (*H_d_*=0.81), and genetic structure was lower (*G_ST_*=0.143) in mainland populations vs peninsular populations (*H_d_*=0.71, *G_ST_*=0.358 respectively). Genetic diversity was negatively associated with elevation but positively with rainfall. Two mainland and one peninsular ancestral haplotypes were reconstructed. Peninsular populations were as isolated among them as with mainland populations. Peninsular haplotypes formed a group with one mainland coastal population, and populations across the gulf shared common haplotypes giving support to regular gene flow across the Gulf. Gene flow is likely mediated by bats, the main pollinators and seed dispersers. Niche modeling suggests that during the Last Glacial Maximum (c. 130 ka), OPC populations shrank to southern locations. Currently, Stenocereus thurberi populations are expanding, and the species is under population divergence despite ongoing gene flow. Ancestral populations are located on the mainland and although vicariant peninsular populations cannot be ruled out, they are likely the result of gene flow across the seemingly formidable barrier of the Gulf of California. Still, unique haplotypes occur in the peninsula and the mainland, and peninsular populations are more structured than these on the mainland.

## INTRODUCTION

The present distribution of species is the consequence of historical processes, ecological interactions, and events of dispersal and/or vicariance. The geographic range of a species can be split into separate discontinuous populations that may eventually lead to allopatric speciation [1, 2]. In addition, there may be associated long-distance dispersal events and founder effects [3]. Major forces behind the actual shaping of the geographic distribution of species include the movement of land masses through the action of tectonics, dispersal distance, changes in climate across the species range, and biotic interactions [4]. Examples of tectonics leading to the formation of land barriers include mountain development through fault blocks and alleviated valleys of the Basin and Range Province, reverse thrust, and the broad consequences of rifting of the continental crust [5]. Events such as the glaciations confined species to refugia where they persisted during the Pleistocene glacial maxima [6, 7, 8], and extant populations in these refugia functioned as reservoirs for recolonization once conditions were suitable for range expansion. Tectonics, climatic shifts, and repeated cycles of glaciation have been proposed as important components in the formation of vicariant populations [9] that might produce speciation or fuse divergent populations as barriers relaxed [10]. As noted by Cavin (2017) [11], vicariance is the most straightforward hypothesis to explain disjointed species distributions. For example, climate and tectonics closely match speciation in Cichlid fish speciation and phylogeny [12]. In linear stream systems, using flow and habitat characteristics Cañedo-Argüelles et al. [13] described three kinds of dispersers, these most affected by local factors, intermediate dispersers responding to landscape factors, and strong dispersers with no major pattern at the regional-scale. The former is the most likely to have disjoined populations. In high-elevation *sky islands* have distinct, closely related species that became separated from a single taxon during glacial cycles [14]. For lower elevation taxa, dispersal better explains species distributions. For insular systems, the explanation of older taxa in newer islands led to the proposal of vicariant metapopulations as an alternative between long-distance biogeographic dispersal [geodispersal] and ecological short-distance species movements [15, 16].

The evolutionary mechanisms of vicariance and colonization through dispersal shape the genetic structure and divergence among populations [1, 2]. However, separating the mechanisms leading to allopatric differentiation is difficult, and represents a major challenge in biogeographical research. Gene flow [17, 18] acts as a major unifying force holding together a common gene pool, and the main forces behind population differentiation are random events like mutation or genetic drift, and natural selection. The latter, drive adaptation, particularly when species have a wide genetic basis, wide distribution range, and are exposed to extensive environmental geographic variability [19]. In any case, for differentiation to occur, reduced gene flow is important. Because of its length (about 1200 km) and narrowness (40-180 km), the peninsula of Baja California, Mexico is in an almost insular environment. Baja California isolation is the result of tectonic processes that separated the peninsula from the mainland by gradually creating new ocean floor during the last 12 million years (late Miocene) [20]. The movements of the tectonic plates of the Pacific and North America resulted in the gradual emergence of the Gulf of California c. 4-5 million years ago as well as the creation of the Rivera plate through what is known as “strike-slip faults” [21, 22]. These geological episodes led to remarkable ecological changes to the biota of the peninsula, to evolutionary processes of allopatric speciation [23–26], and reduced gene flow [i. e., 27].

Although the Gulf of California, is not very wide (about 200 km at its widest point and much less among islands of the midriff region), is recognized as a geographical barrier for many species of the Mexican Pacific coast [8, 28]. Plant and animal species derived from ancient populations living before and during the opening of the Gulf probably evolved in geographic isolation under new ecological, geological, and oceanographic determinants. These factors led to a high percentage of endemism in the peninsula and the gulf islands [25, 29]. Also, Wilder et al. [30] showed that although small islands have unique community assemblages, larger islands are strongly related to the continental sources.

As a result of the peninsular isolation many species show morphological, genetic, or functional differences with mainland species [29]. Within the peninsula, some species have been further isolated by the complex topography of its mountain ranges, the active tectonics, and sea level changes that at times turned what is now the peninsula into archipelagos and biological islands [24–26, 31]. Evolutionary processes including vicariant differentiation as well as recent events of dispersal have played a key role in population differentiation [32, 33]. A striking example of population genetic differentiation across the Gulf of California, are boojum trees (*Fouquieria columnaris*) showing two distinct clades: a homogeneous peninsular clade and a highly diverse mainland clade (34).

Columnar cacti are prominent elements of the warm drylands of North America. Some species, like the saguaro cactus (*Carnegiea gigantea* (Britton & Rose) Engelm.), or the sahuira (*Stenocereus montanus* (Britton & Rose) Buxb.) are only present in the mainland, either disappeared in the peninsula or never dispersed across the Gulf of California [35]. Other species are only found in the peninsula such as the cochal and the creeping devil [*Myrtillocactus cochal* (Orcutt) Britton & Rose, *Stenocereus eruca* (Brandegee) A.C. Gibson & K.E. Horak, respectively) Their presence are likely the result of vicariant events. Still other columnar cactus species such as the cardón sahueso and the pitaya agria (*Pachycereus pringlei*, (S. Watson) Britton & Rose, *Stenocereus gummosus* (Engelm.) A.C. Gibson & K.E. Horak, respectively) are found in the peninsula and the coastal mainland. Aside from the senita (*Lophocereus schottii* (Engelm.) Britton & Rose), that has a highly specialized pollinator mutualism by moths [32], the organ pipe cactus (OPC, *Stenocereus thurberi* (Engelm.) Buxb.) is the only species of columnar cactus having an extensive distribution both in the mainland and the peninsula. The OPC differs from the senita in being pollinated and dispersed by long-distance flying vertebrates, in particular the nectar feeding bat *Leptonycteris yerbabuenae* and several species of perching birds [35]. This feature makes it a suitable model to test the relative contributions of vicariance and geodispersal in its evolution, and the evolution of columnar cacti - bat mutualisms.

The variation, structure, and gene flow of mainland populations of *S. thurberi* were studied using nuclear genetic markers by Bustamante et al. (2016) [37]. They found moderate genetic structure because of the strong gene flow among the populations. The present study focused on elucidating the relative contributions of vicariance and geodispersal in the biogeographic context of the OPC. To do so, we analyzed the phylogeographic pattern, the genetic diversity and structure, and the historical and ecological relationships of twelve populations, comparing the mainland and Baja Californian populations.

## MATERIAL AND METHODS

### Study plant

*Stenocereus thurberi*, also known as organ pipe cactus (OPC) and the pitaya dulce, is a multi-stemmed 3-8 m tall columnar cactus of the arid regions of northwestern Mexico [28]. As most large columnar cacti, it is a key element of the desert dynamics providing shelter and food to many organisms [35]. Also, it is of ethnobotanical interest, as its fruits are staple food during the driest and hottest part of the year, and their woody remains are used as building materials by native people [35]. It has single, perfect, whitish flowers with nocturnal anthesis [36]. Flowers open shortly after dusk and close during the morning of the next day. *Stenocereus thurberi* usually starts flowering from late April to mid-May, and depending on the geographic location, blooming lasts 8 to 16 weeks [36–38]. During this time, its reproductive success depends primarily on bat pollination and secondarily on hummingbirds, perching birds and hawkmoths [36]. Later in the season, nectar feeding bats and perching birds disperse their seeds [39].

### Sampling

Stem tissue from 20 to 25 well-separated adult individuals per population was collected from 12 natural populations of *S. thurberi*. Seven populations were sampled in Sonora, Mexico (mainland group, MAIN from now on) covering the mainland distribution range, and five populations in the peninsula of Baja California (peninsular group, PEN from now on), ranging from the southernmost population near Los Cabos to the Vizcaino region (Fig 1; S1 Appendix). Plant material collected in the field from MAIN populations was stored in an ice chest in the field and frozen at the end of the day. Peninsular samples were dry stored in silica gel after field collection. Once in the lab, all samples were stored at -80 °C.

**Figure 1.**
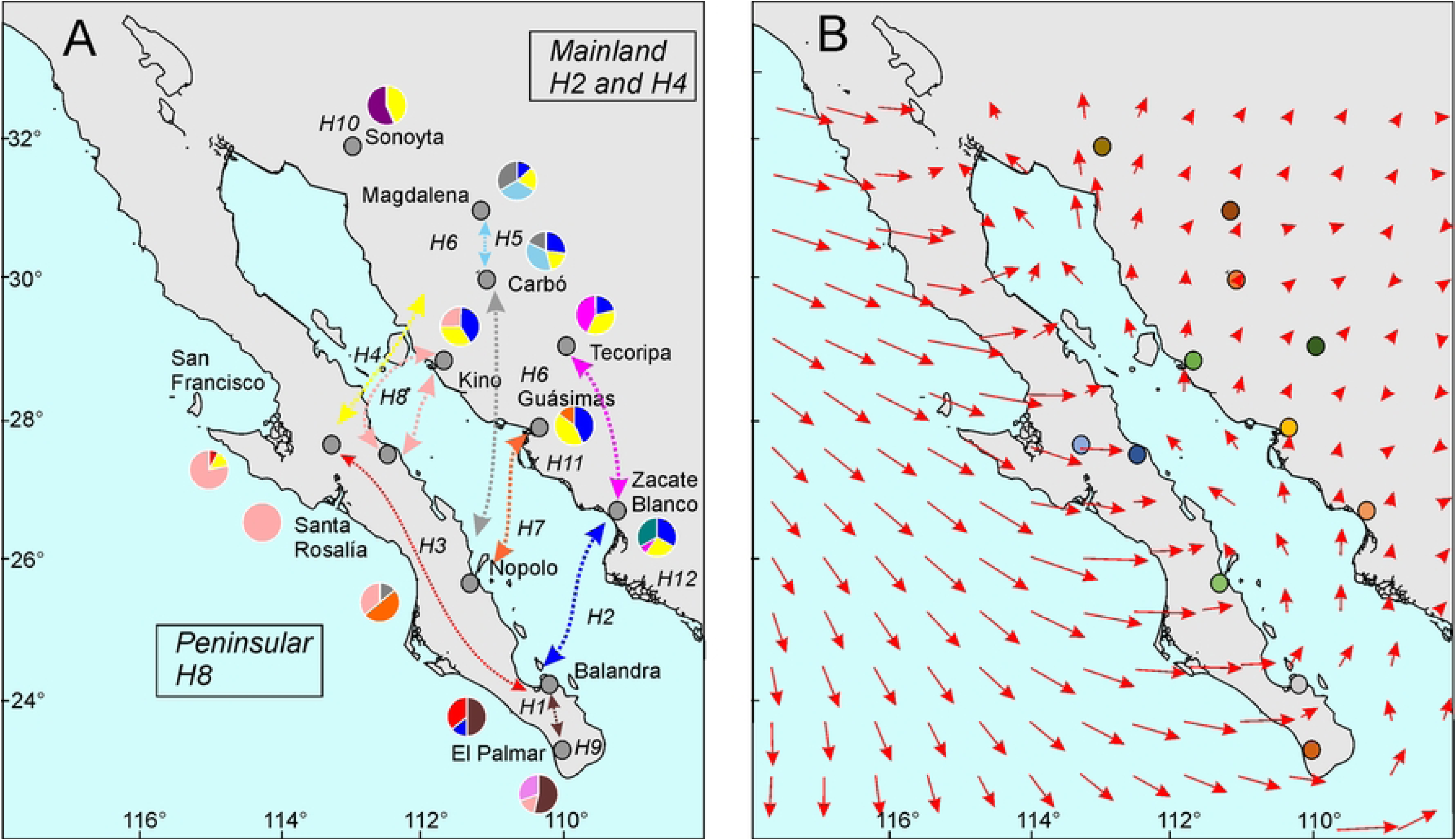
Distribution of the organ pipe cactus [green shading; modified from Turner et al. [27] and populations studied [blue stars, blue font]. Major nature reserves are indicated by italic names. Major cities in grey circles.

### DNA extraction, PCR amplification, and sequence alignment

Total genomic DNA extractions were conducted using a modified cetyltrimethylammonium bromide (CTAB) protocol [for details see 37]. We obtained the total DNA from a set of 15-20 individuals per population. Three non-coding chloroplast fragments (two polymorphic intergenic sequences: rpl32-trnL and trnL-trnF [40] and petB intron D4 gene [40]) were PCR amplified and sequenced from a total of 216 individuals (see S2 Appendix for the full PCR procedure).

Multiple alignments of the sequences were obtained with MUSCLE [42] for a total of 176 samples, and the regions were concatenated with DnaSP v6 [43]. All sequences were deposited in the NCBI GenBank (rpl32-trnL: KU59180-KU59339, trnL-trnF: KU59340-KU59485 and PetB intron D4 gene: KU31391-KU1554).

### Genetic diversity, haplotype phylogeny and network

Diversity indices were calculated for each population and geographic group (i. e., MAIN group and PEN group) as measures of genetic variation for cpDNA. Using DnaSP v6 [43] we estimated the number of segregating sites (*S*), of observed number of haplotype (*h*), average nucleotide diversity (π), average haplotype diversity [*Hd*; 42]. The significance of these tests was evaluated using the coalescent simulation implemented in DnaSP with 10,000 permutations.

Unique haplotypes were selected from a concatenated dataset for phylogenetic analysis with Bayesian inference (BI). We determined the best fit models of nucleotide substitution using jModeltest v0.1.1 [45] under Akaike Information Criterion [AIC]. Bayesian analysis was performed with MrBayes v3.2.1 [46] under the following partition: trnL-rpL32 (TN93+G; AIC = 6799.8; lnL = -3223.6); trnL-trnF (GTR+G; AIC = 6302.2; lnL = -3119.1); PetB intron D4 gene (TN93+I; AIC = 2719.4; lnL = -999.7). Chloroplast sequences of *S. gummosus* were downloaded from Arias et al. (2005) [47], Hernández-Hernández et al. (2011) [48] and Plume et al. (2013) [49] and used as an outgroup to root the phylogenetic tree (see S2 Appendix for the full Bayesian inference). We implemented a median-joining [50] method to build an unrooted haplotype network using coalescent simulations in Network v.10.2.0.0 (available on www.fluxus-engineering.com, 2020). The intra-specific relationships between haplotypes were analyzed using the least cost criterion, treating gaps as single evolutionary events and indels as a fifth state of character [51].

### Genetic Structure

The observed genetic variation among and within populations and among PEN and MAIN groups was partitioned by a hierarchical analysis of molecular variance (AMOVA) using ARLEQUIN v.3.5.1.2 [52]. Three hierarchical divisions based on the genetic variance were identified: a) within populations, b) among populations within groups, and c) among groups using a non-parametric permutation procedure incorporating 10,000 iterations. Pairwise *F*_ST_ [53] were calculated using ARLEQUIN v.3.5.1.2. Then, a Mantel test was performed to assess the correlation between the genetic distance *F*_ST_ / (1 - *F*_ST_) matrix and the geoid distances (km) matrix. In addition, estimates of *G*_ST_ structuring were calculated for each of the two population groups and for the whole sample. To determine the presence and hierarchy of barriers to gene flow occurring among populations, we analyzed the Nei’s matrix of genetic distances with the Monmonier maximum difference algorithm [54]. BARRIER 2.2 was used to determine hypothetical barriers within the distribution of populations.

To study the genetic structure of *S. thurberi,* we used the model-based Bayesian clustering method of STRUCTURE v2.3.4 [55].

### Coalescent inferences of gene flow and divergence timing

Using an isolation with migration (IM) model implemented in IMa v2.0 [56], we estimated the effective population sizes (*N_e_*) of each population group, and the gene flow between the PEN and MAIN populations using the relative rates of population migration (*M*) between groups to capture the population dynamics of the population groups in the early stages of differentiation [57], This model uses a Bayesian coalescent-based Markov chain Monte Carlo (MCMC) method to estimate the posterior probability density of parameters [58]. The model involves several simplifying assumptions, such as neutrality and non-recombination within loci, lack of genetic contribution of unsampled populations, and random mating in ancestral and descendant populations [56]. Preliminary runs were performed to assess the convergence of the MCMC chains on the data stationary distribution and optimize upper bounds on prior distributions (*q* = 30, *t* = 5, *m* = 50; where *q* = population size, *t* = divergence time, and *m* = migration rate) and to optimize heating schemes. Final analyses consisted of three runs of 50 geometrically heated chains with burn-in of 5 x 10^6^ steps. The heating scheme used a geometric model with parameters *h_a_* = 0.9 and *h_b_* = 0.5. A total of 1,000,000 genealogies were saved after the three long runs and used to calculate parameter values and likelihood ratio tests of models [59]. Demographic parameters were scaled at generation time and neutral mutation rate. We used a generation time of twenty years. In the absence of a well-calibrated estimate for the *Stenocereus* genus, we applied a generic mean mutation rate (μ) of 7 × 10^-9^ base substitution per site per generation for the cpDNA [60], and the average geometric mutation rate of the markers was used to scale the outputs to demographic units.

We used BEAST v2 [61] to produce a calibrated tree over time sequenced regions of cpDNA. This analysis was performed using the best evolution molecular model based on jModelTest 2.1.4 (see results), to determine the approximate dates of the divergence events between the haplotypes of the population groups of *S. thurberi*, i. e., via vicariance (> 2.5 million years ago) or dispersal (<2.5 Ma) using a relaxed clock with a log-normal model and the Yule (pure-birth) speciation process as the tree prior [62]. To calibrate the tree in years, we used one calibration point estimated by Hernández-Hernández et al. (2014) [63]. A posterior distribution of time-calibrated trees was estimated in a 100-million generation MCMC run, with samples taken every 1000 generations. The resulting trees were summarized using TREEANNOTATOR from the BEAST package of R.

### Ecological Niche Model (ENM)

To model the ecological niche (ENM) and to reconstruct the potential geographic distribution of *S. thurberi* in different historical periods we used Maxent v3.3 [64]. A total of 252 records covering the species distribution range were used. These included the populations of the present study, the geo-referenced populations in Bustamante et al. [37], and the records of the GBIF database (http://www.gbif.org/). To avoid redundancy and overfitting, we removed repeated occurrences. We used 19 bioclimatic layers with a spatial resolution of 2.5 arc-minute (WorldClim database v1.4) [65] to describe the general conditions of the area at the different modeled times. Highly correlated variables were excluded (Pearson’s *r* ≥ 0.7), variables with the larger contribution to model development and likely more biological importance were retained (S3 Appendix). As a result, climatic conditions were measured as a function of 9 bioclimatic variables. These bioclim variables were used to identify suitable actual climatic areas (1960–1990), possible refuges in the paleodistribution of last glacial maximum (LGM, *c*. 21 ka; climate system model 4, CCSM4), and suitable environments during the last interglacial (LIG, *c*. 120-140 ka; climate system mode NCAR-CCSM). We assumed that climatic preferences did not change over time. Model validation was performed using the default Maxent v3.3. [64] configuration with 10 independent subsample replicas.

To determine the role of environmental variation across the distribution of *S. thurberi* on the geographic patterns of retained genetic diversity, we performed a multiple linear regression using SPSS 20 (SPSS Statistics for Windows, Version 20. Armonk, NY: IBM Corp.). We used the 12 populations as replicates, the averaged genetic diversities (*Hd* and π) calculated in DNAsp as response variables, and the most important variables from the ENM as explanatory variables. Models were tested for the pooled populations and separately for the MAIN and PEN populations.

## RESULTS

### Chloroplast data, haplotype phylogeny and network

The three analyzed chloroplast non-coding regions gave rise to sequences spanning 2338 bp. We found 10 segregating sites with 12 different haplotypes (H1 to H12) for the 12 populations. The information of the estimated genetic variation is summarized in Table 1. Nucleotide diversity (π) ranged from zero to moderate (0.9×10^-3^), and haplotype diversity (*Hd*) from zero to 0.76. The lowest diversity was found in northern Baja Californian populations. Pooled haplotype diversity and average population diversity in the mainland (0.81 and 0.68, respectively) was higher than peninsular haplotype diversity (0.72 and 0.47, respectively, Table 1).

**Table 1.**
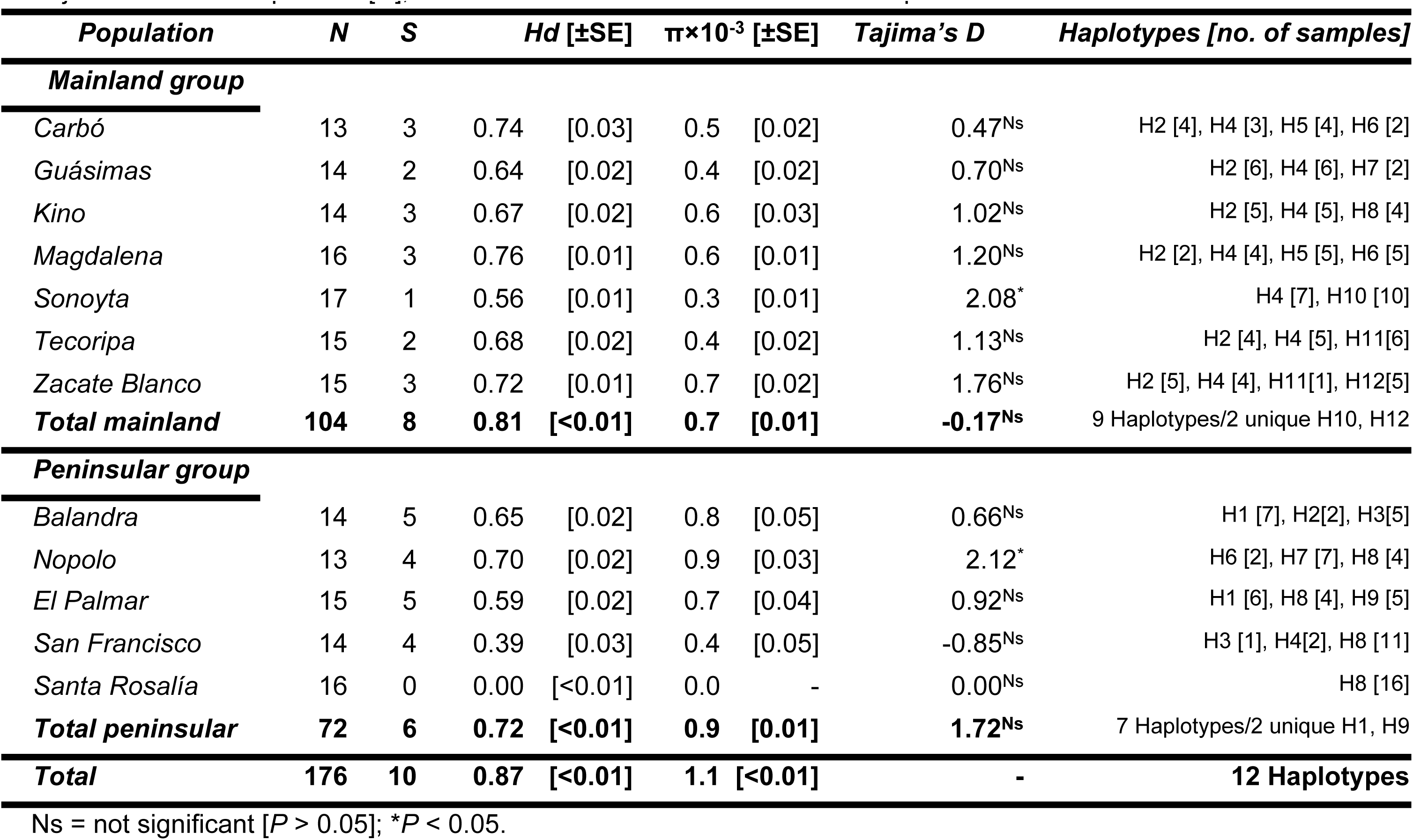
Number of polymorphic sites [S], haplotype diversity [Hd], nucleotide diversity [π], and Tajima’ s D test of population expansion/contraction in populations of Stenocereus thurberi sampled in the mainland and the peninsula of Baja California. Sample size [N], Hd and π ± standard error of the mean in parentheses.

*Tajima’s D* tests showed that most populations are neutral, without indications of selection or population changes. Only two populations (Sonoyta and Nopolo) displayed a slight but significant deviation from neutrality (Table 1). There was no association between haplotype diversity and topographic variables. However, when MAIN and PEN groups were analyzed separately, a significant negative association with elevation was found for the PEN group but not for the MAIN group (S4 Appendix). Also, a highly significant relationship between *Hd* and annual rainfall showed that *Hd* rapidly increased with rainfall to reach a plateau at about 400 mm annual rainfall (S4 Appendix) A phylogenetic reconstruction estimated by Bayesian analysis partially resolved the haplotype relationships and revealed two main clades: one comprising the mainland haplotypes H1, H2, H3, H5, H6, H8, and H9, and the other peninsular haplotypes H11 and H12 (Fig 2A). Both clades are well-supported (posterior probabilities of 0.75 and 0.96, respectively). Within the large group, some divergent haplotypes also segregated into a mostly Baja Californian clade (H1, H3, and H8) with high support (0.99), and the remaining haplotypes included mainland and peninsular haplotypes. The remaining haplotypes (H4, H7, and H10) clustered with all the above forming a monophyletic group (posterior probability of 1.00), but their phylogenetic relationships were not resolved (Fig 2A).

**Figure 2.**
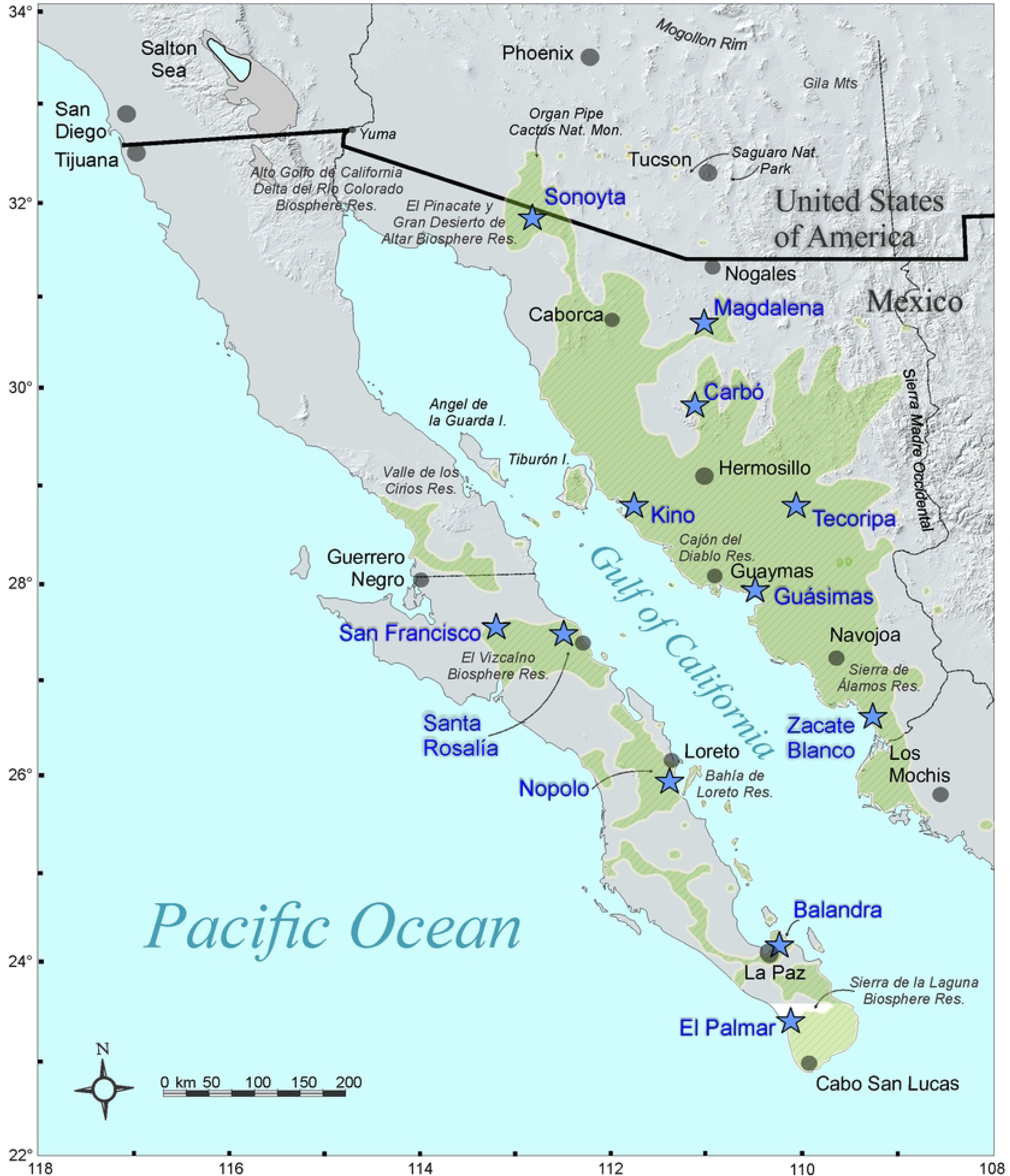
Bayesian analysis of 12 cpDNA haplotypes of *Stenocereus thurberi* based on three cpDNA regions [A]. Posterior probabilities are indicated in the branches of the tree. Minimum spanning network of genealogical relationships among haplotypes [B]. The size of circles corresponds to the frequency of each haplotype and the small black circles are mutational steps. Phylogenetic tree under a relaxed clock model [C]. The estimated times of divergence of nodes are indicated. The blue error bars at the nodes represent the 95% confidence limits.

Additionally, the median joining network of 12 haplotypes (Fig 2B) revealed partial divergence, with only one ambiguity between the haplotypes H6 and H7, pointing to gene migration among MAIN and PEN populations. The PEN population of Nopolo shares its three haplotypes with MAIN populations, in particular with the two coastal populations across the Gulf of California (Table 1, Fig 2B, S5 Appendix). In all cases, the most frequent haplotypes (H2, H4 and H8) were in the center of the network. The least common haplotypes were located at the tips of the network, in the southernmost populations of both the peninsula (H9, El Palmar) and the mainland (H12, Zacate Blanco). Four haplotypes were found only in one population each: two in the mainland at the northern (H10, Sonoyta) and southern (H12, Zacate Blanco) latitudinal distribution of MAIN populations (Table 1), and two in the Cape Region at the southernmost tip of the peninsula (H9 El Palmar and H2 Balandra).

Haplotypes H5, H10, H11, and H12 were only found in the MAIN populations. The northern PEN population of Santa Rosalía had only one haplotype (H8; Table 1), while haplotypes H1 and H9 were only found in the southernmost populations of the Cape Region of the peninsula. Haplotype H7 was the only one that was shared by one coastal MAIN and one PEN population (Fig 2B).

The network showed three main lineages or haplogroups, each derived from a common haplotype: [1] The haplogroup that comprises the H2 and the uncommon haplotypes derived from H2 (H5, H6 and H9), with a wide geographic distribution in the mainland [except in Sonoyta, the northernmost mainland population] and marginally in the peninsula. [2] The H4 haplogroup, including H7, H10, H11 and H12, all of them widely distributed throughout the mainland and almost absent in the peninsula. [3] The H8 haplogroup, also comprising the H1 and H3, found throughout the PEN region and only in the MAIN population of Kino (Figure 2B, S5 Appendix).

The most probable tree chronogram (Figure 2C) indicated that *S. thurberi* originated *ca.* 2.05 Ma (95% HPD, 1.06 - 3.23 Ma) during the late Gelasian of the Pleistocene. The largest haplogroup originated about 1.56 Ma (95% HPD, 1.16 - 2.24 Ma), when the Gulf of California was already well established. It includes three clades, one composed of H6, H7 and H9 haplotypes, diverging roughly 0.69 Ma (95% HPD, 0.009 - 1.52 Ma), located in the mainland and the peninsula. Two additional haplogroups diverged about 1.21 Ma (95% HPD, 1.01 - 1.67 Ma). The first, including H2 and H5 haplotypes, diverging ca. 0.56 Ma (95% HPD, 0.01 - 1.63 Ma), mainly present on the mainland, and the other distributed along the peninsula and branching back 0.65 Ma (95% HPD, 0.03 - 1.38 Ma). The smallest haplogroup was composed of the most continental-distributed haplotypes (H4) and other mainland-restricted haplotypes (H10, H11 and H12) at 0.89 Ma (95% HPD, 0.001 - 2.04 Ma).

### Genetic differentiation and population structure

The STRUCTURE analysis including all populations and using *K* = 2 confirmed two geographic genetic clusters, one in the mainland and one in the peninsula (Fig 3A). Some individuals had an inconsistent genetic signature indicating genetic exchange between MAIN and PEN populations, particularly those on the central Gulf of California coast. An analysis with *K* = 3 separated the two main population groups, and the northernmost MAIN population (Sonoyta; S6 Appendix). With *K* = 4 there was an additional, well-defined cluster separating the populations of the Cape Region on the southern tip of the peninsula (Balandra and El Palmar; S6 Appendix).

**Figure 3.**
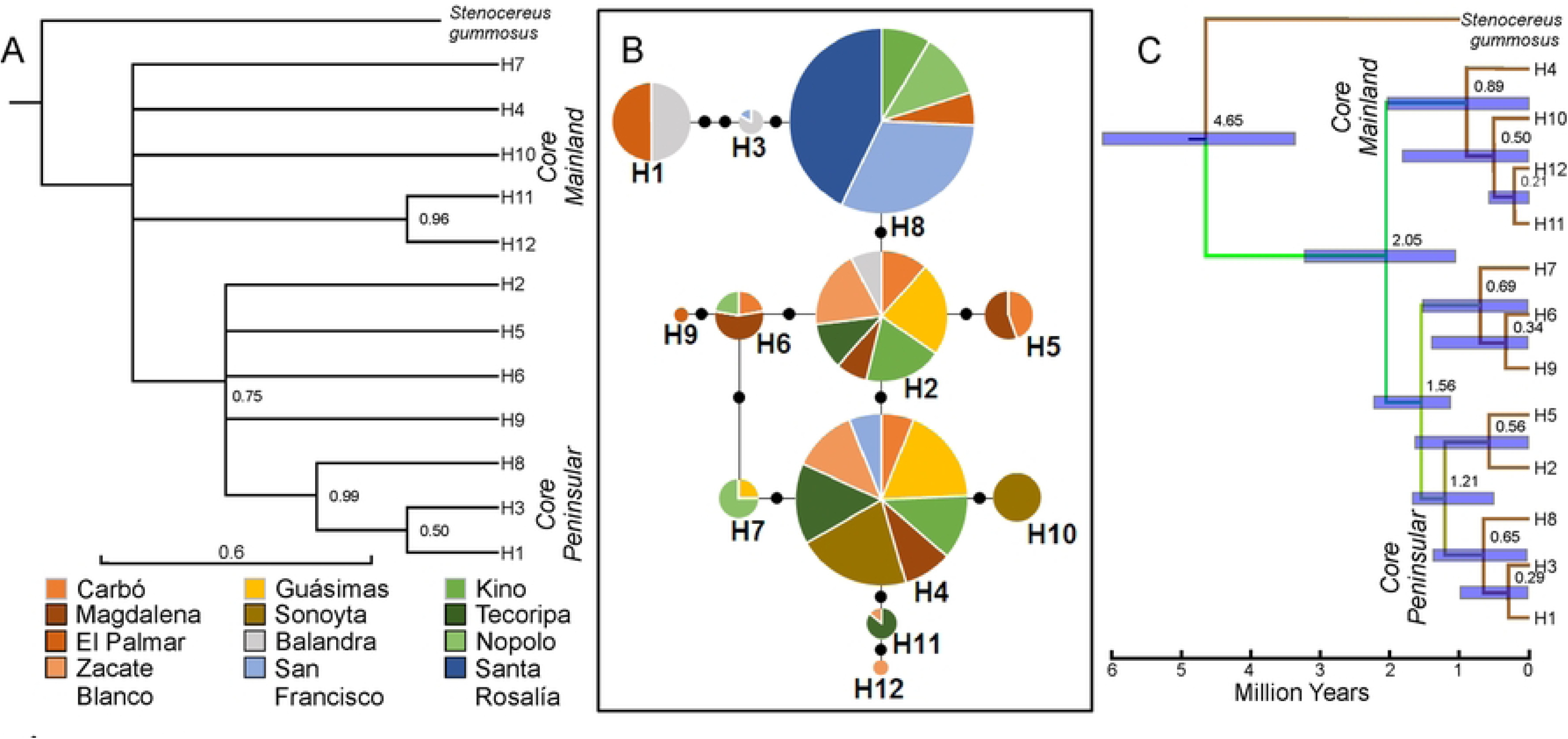
STRUCTURE assignment of the 176 samples according to geographic groups for *K* = 2 [using admixtures and correlated allele frequencies], for samples from the mainland [left, *K* = 4], and samples from the peninsula [right, *K* = 3] [A]. PCA ordination of the individuals of the mainland [B], and peninsular geographic region according to their genetic structure [C] on the first two principal components [percentage variance explained in parentheses].

The STRUCTURE analyses conducted only on the PEN populations revealed that the most likely number of clusters according to the Evanno et al. (2005) [64] test was *K* = 3. One cluster was found throughout the peninsula (Figure 3A, grey), another included only the monomorphic population of Santa Rosalía (yellow), and a third clustered the two Cape Region populations (Balandra and El Palmar, light purple). For the mainland, the most likely number of clusters was *K* = 4: one including widely distributed individuals (red), a second cluster with plants only from Kino (blue), a third group including thornscrub plants from Tecoripa and Zacate Blanco (purple), and a cluster dominating in the northernmost population of Sonoyta (green). These individual assignment patterns were consistent with the genealogy (Fig 3B).

The subgroups detected by the PCA in the MAIN populations differentiated the coastal populations, the inland populations (with higher values along PC2), and the northern population of Sonoyta (Fig 3B). Also, in the PEN populations three subgroups emerged: the Cape Region (Balandra and El Palmar), central (Nopolo), and northern (San Francisco and Santa Rosalía) (Fig 3C). Consistent with the results above, an AMOVA analysis showed significant genetic differences between geographic groups (*F*_CT_ =0.443**), between populations within groups (*F*_SC_=0.367**), and within populations (*F*_ST_=0.648**). The difference between the two groups explained 44.4% of the total variation of cpDNA while 35.2% was attributed to differences between samples within populations; and the remaining (20.5%) was attributed to differences between populations within groups. Total genetic differentiation (0.64) indicates a high genetic structure (S7 Appendix). On the other hand, the *G*_ST_ value was moderate for the MAIN populations (0.143), and higher for the PEN populations (0.358) and for the whole sample (0.313).

The Mantel correlation between genetic and geographic distance matrices detected significant correlation for the set of populations (*R*^2^= 0.22; *P* =0.003; n_m_=66) and for the PEN populations (*R*^2^= 0.35; *P* =0.04; n_m_=10), but it was near the limit of statistic acceptance for the MAIN populations alone (*R*^2^= 0.21; *P* = 0.07; n_m_=21) (Fig 4A; S8 Appendix).

**Figure 4.**
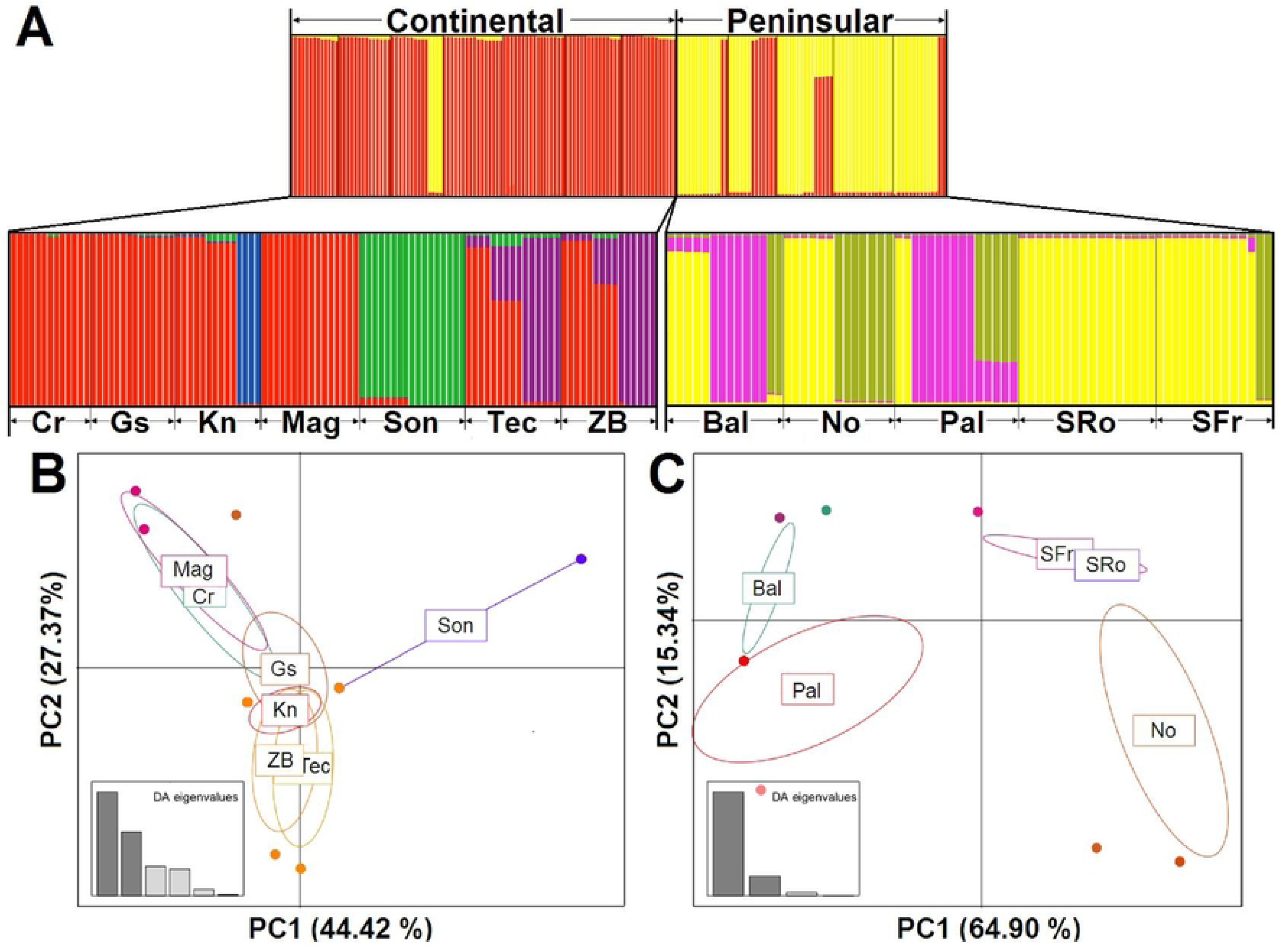
Correlation between the paired geographic and genetic distances for all [green], only MAIN [blue] and only PEN [red] populations of *S. thurberi* [A]. Estimation of geographic barriers to gene flow among populations using the genetic and geographic distances [B]. Delaunay triangulation of geographic distances [green], Voronoi tessellation showing the domain of each population [magenta]. The thickness of each edge [red] and the number in the barriers is the percentage of times it was included in one of the 200 bootstrap repetitions to compute barriers. All other values in color magenta are less than 10.

Using the Monmonier maximum difference algorithm on the Nei’s genetic distance matrix we found that 1) mainland populations had little isolation in terms of genetic barriers (Fig 4B). The exception, with bootstrap values just above 50%, was the northernmost mainland population of Sonoyta (Fig 4B). 2) The Gulf of California is the most important barrier to gene flow. However, within the peninsula, the northern (San Francisco and Santa Rosalía) and the southern PEN populations in the Cape Region were as isolated among them as populations on the mainland and the peninsula. The northern and southern PEN populations are separated from the rest of populations by the Gulf barrier with a bootstrap support of 100%. The central PEN population of Nopolo showed a higher connectivity across the gulf than to neighboring PEN populations to the north and south (Fig 4B).

### Coalescent analysis of divergence history

Relative population migration rates (*M*) between groups were estimated using IMa for the MAIN and PEN populations, reaching convergence with high values of effective sample sizes in all parameters (ESS>1100). IMa estimated approximate *N_e_* for the MAIN group as 3.8 x 10^5^ individuals (90% HPD = 1.6 x 10^5^–1.4 x 10^6^) and *N_e_* for the PEN group as 2.7 x 10^5^ individuals (90% HPD = 1.6 x 10^5^–1.6 x 10^6^), making the mainland *N_e_*slightly larger (S9 Appendix), while ancestral *N_e_* was about 3.8 x 10^5^ individuals (90% HPD = 1.7 x 10^5^–1.1 x 10^6^). The estimated migration between the two population groups was higher than one migrant per generation (S9 Appendix). The effective number of migratory events per generation from the peninsula to the mainland (PEN → MAIN), 2*NM*, was 2.3 (95% HPD 0.74–7.2 migrants per group per generation), while migratory events from the mainland to the peninsula (MAIN → PEN) were estimated at 1.3 (95% HPD 0.24–5.8 migrants per group per generation). Both estimates broadly overlap, suggesting bidirectional gene flow across the Gulf of California.

### Ecological niche profiles

The estimated ecological niche models (ENM) predicted the current distribution of *S. thurberi* (Fig 5) with high certainty (AUC values >0.97 for training and testing data). Jackknife tests of the importance of variables revealed that precipitation of the driest quarter, precipitation of the coldest quarter, and annual precipitation were the top three ranked variables when used in isolation. The simulation of the three periods, current, Last Glacial Maximum (LGM, 21 Ka, MIROC model), and Last Interglacial (LIG, Sangamonian stage, 120-100 Ka) times indicated extensive changes in the distribution of *S. thurberi*.

**Figure 5.**
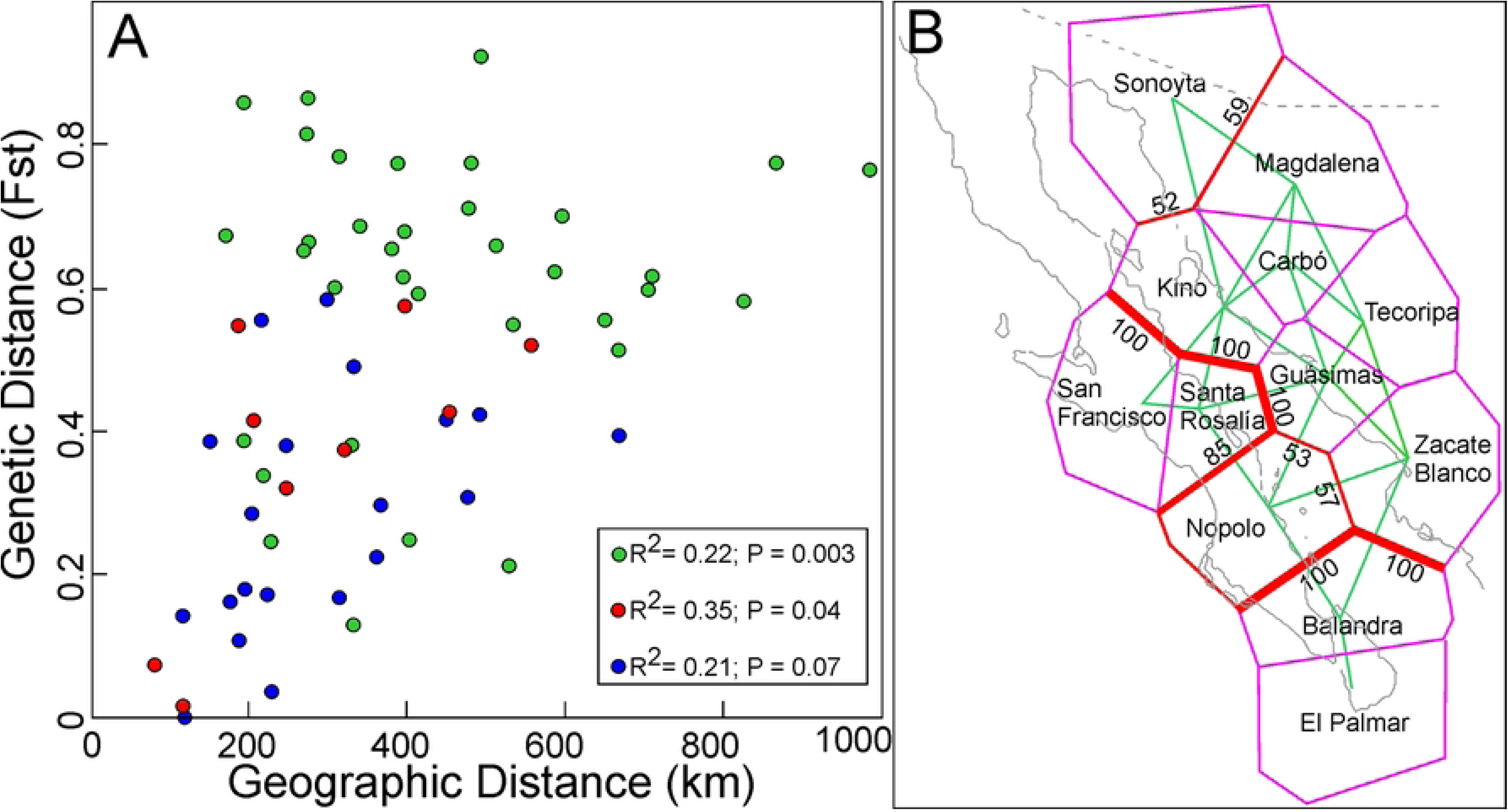
Predicted environmental suitability for *Stenocereus thurberi*, using ecological niche models [ENM]. ENM results are shown for: A] the current, B] the last glacial maximum [LGM, 21 Ka, MIROC-ESM model], and C] the last interglacial period [LIG, 120-100 Ka] time periods.

The predicted climatically suitable habitat for the species coincided with the current geographic distribution of the species (Fig 5A; [28, 37]), suggesting that the suitable habitat available for the species was substantially reduced during the LGM. Apparently, refugia were in coastal areas in the southwestern Sonoran Desert. This location can be explained by the lower sea levels, maritime climate, and milder temperatures in southward locations where Las Guásimas, Zacate Blanco, El Palmar, Nopolo and Balandra populations are today (Fig 5B). These analyses show a potential range shift affecting the distribution through time, increasing suitable habitat for the species towards the present.

Under the modeled climatic conditions of the LIG period, the climatically suitable habitat for *S. thurberi* was present in both areas. In the mainland, a suitable patch included areas where the Kino and Guásimas populations are currently found. The second, much larger patch, included the populations of the north-central peninsula (San Francisco, Santa Rosalía and Nopolo; Figure 5C).

Stepwise multiple linear regressions of the average genetic diversities (*Hd* and π) and the main climatic parameters identified in the PCA for the ENMs showed that seasonality and mean annual temperature were significantly correlated with π, while Hd was only related to annual precipitation (S10 Appendix). When doing the analysis separately for PEN populations, annual precipitation was the most important factor determining π and *Hd*. In the case of *Hd*, mean annual temperature also contributed with significant added variance (S10 Appendix). In the mainland annual rainfall was not significant when using linear regressions, and isothermality and mean annual temperature explained a significantly large proportion of the variance in *Hd* (S10 Appendix). Simple, non-linear models explain better the relationship with annual precipitation. In particular, the variation in *Hd* is better explained by the annual precipitation using an inverse function. Annual precipitation is not linearly related to genetic diversity, but a non-linear inverse fit shows that precipitation is a very good predictor of π and *Hd* across populations (S10 Appendix, S3 Appendix).

## DISCUSSION

Different models have been proposed to explain the evolutionary processes producing the diversity and endemism of flora and fauna of the Baja California peninsula and the many islands product of the tectonic forces shaping the evolution of the gulf [25, 33, 67]. In the case of Baja California, there is agreement that at least three major fragmentation processes associated with the creation of the Upper Gulf of California occurred since the Late Pliocene. Such processes also created the Isthmus of La Paz, which was covered by water during this same period, and the central region of Baja California, which before the Middle Pleistocene was either an island separated from the northern reaches by a shallow canal or a narrow isthmus northward [26, 32]. These processes have been ascribed as the cause for the numerous cases of endemism [29; 68; 69]. Many of them are undoubtedly the product of vicariance [8, 69]. Among these, is the remarkable case of mammilloid cacti where vicariance lead to sister species across the Gulf [71]. However, there are plant species on both coasts (and islands) like *Fouquieria columnaris* [72], *Pachycereus pringlei*, and many others that exhibit little or limited morphological differentiation, despite the geographic barrier of the Gulf of California. This similarity may indicate: 1) that genetic separation might be high but is not apparent at a morphological level (cryptic vicariance; [25]), or 2) that for some species, the Gulf of California is only a partial barrier for gene flow [33]. Any vicariance model of ancestral biota by allopatric isolation during the late Neogene would be the product of population differentiation between species on both coasts of the Gulf, including reciprocal monophily [73–75]. In these cases, the divergence would be the result of geographical barriers, but it would be difficult to exclude recent dispersal [74, 76], and gene flow. *Stenocereus thurberi* is exposed to a variety of biotic and abiotic pressures, including seedling predation [38], drought and temperature extremes, and large disturbances from cyclonic winds and rains. Taken together, the results from *Tajima’s D* test and the historical environmental fluctuations indicate apparent selective processes in Nopolo which is a potential Pleistocene refuge, and in Sonoyta, the northernmost, and probably one of the younger populations. All other populations do not show significant departures from neutrality. Spatially variable selection has been reported in other Neotropical species, the results for *S. thurberi* point in a similar direction [77]. Still, evidence of local adaptation needs to be assessed and the intensity and direction of selection quantified, requiring comparison with phenotypic traits and assessment of survival over time [78]. We did not find higher levels of genetic diversity in what were the likely Pleistocene refuges as found by Sanderson et al. [79] for saguaro with many more markers. Expanding studies to other populations and using information from the whole-genome throughout the species’ natural range should be of interest (particularly those on islands) and reveal adaptive differentiation among populations [4, 80]. Of particular interest is the association of *Hd* and π with precipitation and temperature showing that *S. thurberi* thrives in sites with a well-defined and more predictable monsoon in the fringes of the Sonoran Desert where thornscrub and tropical dry forests develop [81–82]. The negative relationship with elevation in the peninsula reflects perhaps the effects of geodispersal from coastal populations in the mainland and the permanence of ancient populations about 125 m below the actual sea level during the last glacial maximum when low-lying plains were extensive along much closer coasts [29]. The genetic diversity described in our study is high compared to other studies using plastid markers. High levels of genetic diversity have been associated with long-lived woody plants, having high population density, long-distance dispersal of seeds and/or pollen, and extensive geographic ranges [83]. The mean genetic diversity on the mainland was slightly higher than that found in the peninsula (*Hd* = 0.813 vs. 0.713, respectively). Population differentiation was found to be remarkable in the peninsula, while the mainland populations showed less structure (*G_st_* = 0.358 vs. 0.142, respectively), apparently because gene flow is higher on the mainland. In the Cape Region, the peculiarly different climatic and vegetation determinants [84] and the remoteness, apparently led to restricted migration and differentiation. In the southern tip of the peninsula, back in 1892, Katharyne Brandegee recognized a new taxon, *Stenocereus thurberi ssp. littoralis* [Brandegee] N. P. Taylor [85], growing on steep coastal bluffs in a narrow area between San Jose del Cabo and Cabo San Lucas. This subspecies is much smaller, has thinner, greyish stems, pink to magenta flowers, and much smaller fruits than typical *Stenocereus thurberi*. These factors add to the idea of restricted gene flow, isolation, and differentiation as pointed out by Gibson (1990) [86] and support the notion that the Cape Region was isolated by the hypothetical flooding of the isthmus of La Paz [25]. At the same time, the differentiation of the northern populations of the peninsula back the view of an isolated area somewhere north of La Paz and south [26] of Santa Rosalía.

The genetic structure of populations is intimately linked to the activities of their mutualistic pollinators and seed dispersers, including local and migratory movements [87]. The role of chiropterophily in structuring or connecting populations of *S. thurberi* seems to be very important. Nectar-feeding bats are a crucial vector for the pollination and seed dispersal of many columnar cacti [88, 89]. *Leptonycteris yerbabuenae*, the main pollinator of this species [35], and other phyllostomid bats, are efficient pollinators and seed dispersers facilitating the exploration and colonization of new areas [89, 90]. Also, several species of perching birds like orioles, grackles, and woodpeckers (but not doves, as shown by [91]) are known to disperse the seeds (Búrquez pers. obs.) but there is not yet a detailed study of frugivory and dispersal of seeds in any of the columnar cacti of NW Mexico and SW USA.

The high genetic diversity and gene flow we detected within the mainland portion of the distribution range of *S. thurberi,* reveals the absence of strong geographic barriers indicating limited genetic differentiation, and reduced isolation by distance in the mainland populations. For example, the neighboring populations of Carbó and Magdalena share haplotype 5, and Tecoripa and Zacate Blanco share haplotype 11, while the least diverse population is the northernmost locality of Sonoyta with only one haplotype (H10), suggesting recent colonization, or ecological limitations to dispersal [92, 93]. Baja Californian populations of *S. thurber*i are probably the result of ancient colonization from the mainland. They have been isolated from the mainland populations for about 2 million years. However, ongoing gene flow between both groups suggests that even when geographically allopatric with the Gulf of California as a barrier to dispersal, extensive gene flow is still happening, and more than one effective migrant is exchanged between subpopulations per generation. That figure is theoretically enough to stop them from drifting to fixation but to stop divergence, higher levels of gene flow are necessary [10].

Migratory bats are known for their long-range and complex migratory routes [94]. For example, the straw-colored fruit bat, has been shown to traverse long distances across fragmented landscapes and to disperse small seeds by endozoocory across distances up to 50 km [94]. In the case of *L. yerbabuenae*, researchers have found evidence of large, migratory populations on the peninsula, suggesting that bats regularly go across the Gulf of California, potentially using the midriff islands as stepping stones [96, 97].

For example, Medellín et al.) [90] marked individuals of *L. yerbabuenae* captured at foraging sites with colors of powder and found that in night time they can overcome distances of 100 km flying from the cave where they roost to their feeding sites, which exceeds all known distances of other phyllostomids or nectarivores in the world. Examining the distribution of genetic diversity across mainland populations of the *L. yerbabuenae* bat colonies, Ramírez [96] found two clades, but little geographic structuring, and recently Arteaga et al. [98] found the same lack of pattern among peninsular colonies suggesting high levels of gene flow mediated by females. Although there are no direct observations of *L. yerbabuenae* actually using the midriff islands as stepping stones, or flying across the Gulf to reach the peninsula, there are records of this species flying from Tiburon island to the mainland in foraging bouts of about 30 km [99]. Lesser long-nosed bats fly up to 40 km to hummingbird feeders in southern Arizona [100]. Record-breaking long-distance one-night flights were documented recently by Goldshtein et al. [101]. They documented round trip flights from their roosts to their foraging grounds of up to 200 km in the Pinacate y Gran Desierto de Altar biosphere reserve. All these data suggest that *L. yerbabuenae* can fly among islands of the Gulf of California to forage and roost, or to go across the Gulf in one bout. However, if there is paucity in the flying distances of bats, there is even more scarcity of documented seed dispersal distances for bats.

Migration of *L. yerbabuenae* bats between the mainland and the peninsula entail crossing the Gulf of California during the foresummer to find floral resources of columnar cacti during the flowering time early in the season when wind patterns follow a NW-SE direction [102] or later in the season at the onset of the monsoon when strong reversal of the normal surface winds occur and wind follows a roughly S-N direction in the lower Gulf (Fig 6). There is ample evidence for migratory birds taking advantage of winds to assist migration [103], and also for bats [104]. Examining the hypothetical relatedness among populations by haplotype presence shows a remarkable similarity to the summer wind patterns, where the northern coastal populations are probably associated with earlier bat migration across the gulf using the midriff islands as stepping stones, and then within the peninsula. The southern populations on both coasts are related by the likely direct flight of lesser long-nosed bats across the gulf. This hypothetical scenario suggest 1) an early bat migration using either the midriff islands to access floral resources, or taking advantage of the strong winds of the foresummer from the northwest to reach the tip of the peninsula, 2) a return of *L. yerbabuenae* to the mainland to continue foraging on a more diverse set of resources taking advantage of the reverse monsoon winds as noted by the late arrival of *L. yerbabuenae* to the Zacate Blanco population by Bustamante et al. [36], and 3) a return to their wintering grounds late in the season once the wind patterns revert to their autumn - winter pattern. This hypothetical scenario explains the relatedness of populations across the gulf and also resolves the larger number of organ pipe cactus migrants from the peninsula to the mainland.

**Figure 6.**
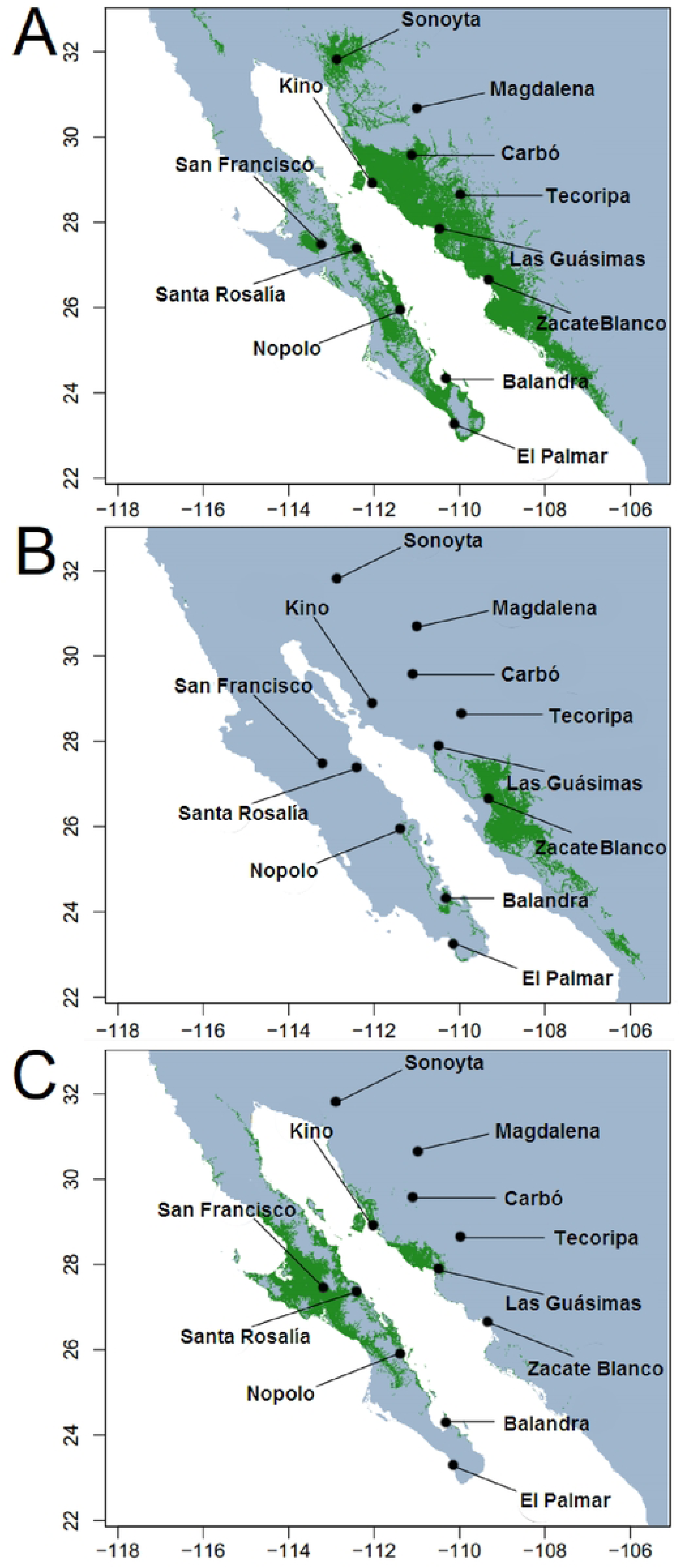
Hypothetical gene flow patterns among populations of *Stenocereus thurberi* across the Gulf of California based on [A] haplotype presence, and [B] surface wind patterns during summertime as modeled by Morales-Acuña et al. [101]. During early autumn, winds return to a generalized pattern like the one on the lower left side of B.

## Acknowledgments

The authors thank Dr. Valeria Souza for her assistance in the field work, and Drs. Erika Aguirre-Planter, Laura Espinosa-Asuar, Santiago Ramírez-Barahona and Josué Barrera-Redondo (all at IE-UNAM) for their valuable help in the laboratory analysis. We also thank Josué Barrera-Redondo, Gustavo Giles and Jonás Aguirre-Liguori for their help with the bioinformatics analyses. Dario Copetti made valuable suggestions to the manuscript. SA thanks Posgrado en Ciencias Biológicas, Universidad Nacional Autónoma de México, and CONACYT for the M.Sc. studentship (480152). This work was also supported by a grant from Dirección General de Asuntos del Personal Académico, UNAM (DGAPA-PAPIIT project: IN213814) to AB.

## Authors Contributions

Conceptualization: Sebastián Arenas, Alberto Búrquez, Luis E. Eguiarte

Field work: Sebastián Arenas, Alberto Búrquez, Enriquena Bustamante, Luis E. Eguiarte

Molecular and bioinformatic procedures: Sebastián Arenas

Formal analysis (analysis and interpretation of the results): Sebastián Arenas, Alberto Búrquez, Enrique Scheinvar

Writing first and final draft: Sebastián Arenas, Alberto Búrquez. All authors reviewed and approved the final manuscript.

## Data availability

Plastid sequences can be accessed through GenBank accessions KU59180-KU59339 for rpl32-trnL, KU59340-KU59485 for trnL-trnF and KU31391-KU1554 for PetB intron D4 [NCBI, https://www.ncbi.nlm.nih.gov/genbank/].

## References

1. Wright S. Genetical Structure of Populations. Nature. 1950; 166, 247–249.

2. Mayr E. Animal Species Evolution. Harvard: Belknap Press; 1963.

3. Dlugosh KM, Parker IM. Founding events in species invasions: genetic variation, adaptive evolution, and the role of multiple introductions. Mol Ecol. 2008;17: 431–449. doi/10.1111/j.1365-294X.2007.03538.x

4. Aguirre-Liguori JA, Ramírez-Barahona S, Tiffin P, Eguiarte LE. Climate change is predicted to disrupt patterns of local adaptation in wild and cultivated maize. Proc R Soc B. 2019;286: 2019048620190486 http://doi.org/10.1098/rspb.2019.0486

5. Lonsdale P. Geology and tectonic history of the Gulf of California. In: Winterer EL, Hussong DM, Decker RW, editors. The Eastern Pacific Ocean and Hawaii. Boulder: Geological Society of America; 1989. doi.org/10.1130/DNAG-GNA-N

6. Bacon CD, Molnar P, Antonelli A, Crawford AJ, Montes C, Vallejo-Pareja MC. Quaternary glaciation and the Great American Biotic Interchange. Geology 2016;44: 375–378. doi:10.1130/G37624.1

7. Bonatelli AS, Pérez MF, Peterson AT, Taylor NP, Zappi DC, Machado MC, et al. Interglacial microrefugia and diversification of a cactus species complex: phylogeography and palaeodistributional reconstructions for *Pilosocereus aurisetus* and allies. Mol Ecol. 2014;23: 3044–3063. doi.org/10.1111/mec.12780

8. Garrick RC, Nason JD, Meadows CA, Dyer RJ. Not just vicariance: phylogeography of a Sonoran Desert euphorb indicates a major role of range expansion along the Baja peninsula. Mol Ecol. 2009;18: 1916–1931. doi.org/10.1111/j.1365-294X.2009.04148.x

9. Nevado B, Contreras-Ortiz N, Hughes C, Filatov DA, Pleistocene glacial cycles drive isolation, gene flow and speciation in the high-elevation Andes. New Phytol. 2018;219: 779–793. doi.org/10.1111/nph.15243

10. Slatkin M. Gene Flow and the geographic structure of natural populations. Science. 1987;236: 787–792. doi: 10.1126/science.3576198

11. Cavin L. Evolutionary patterns in freshwater fishes. *In*: Cavin L editor. Freshwater Fishes: 250 million years of evolutionary history. The Hague: Elsevier; 2017 pp. 127–140. doi.org/10.1016/B978-1-78548-138-3.50005-4.

12. Ivory SJ, Blome MW, King JW, McGlue MM, Cole JE, Cohen AS. Environmental change explains cichlid adaptive radiation at Lake Malawi over the past 1.2 million years. Proc Nat Acad Sci. 2016;113: 11895–11900. doi: 10.1073/pnas.1611028113

13. Cañedo-Argüelles M, Boersma KS, Bogan MT, Olden JD, Phillipsen I, Schriever TA, Lytle, DA. Dispersal strength determines meta-community structure in a dendritic riverine network. J. Biogeogr. 2015;42: 778–790. https://doi.org/10.1111/jbi.12457

14. Arriaga-Jiménez A, Kohlmann B, Vázquez-Selem L, Umaña Y, Rös M. Past and future sky-island dynamics of tropical mountains: A model for two *Geotrupes* (Coleoptera: Geotrupidae) species in Oaxaca, Mexico. The Holocene 2020;30: 1462–1470. doi.org/10.1177/0959683620932977

15. Heads M. Recent advances in New Caledonian biogeography. Biol Rev. 2019;94: 957–980. doi.org/10.1111/brv.12485

16. Hembry DH, Balukjian B. Molecular phylogeography of the Society Islands (Tahiti; South Pacific) reveals departures from hotspot archipelago models. J Biogeogr. 2016;43: 1372–1387. doi:10.1111/jbi.12723.

17. Templeton AR. Human Population Genetics and Genomics. Academic Press. 2018. eBook ISBN. 2018: 9780123860262

18. Aguilée R, Gascuel F, Lambert A, Ferriere R. Clade diversification dynamics and the biotic and abiotic controls of speciation and extinction rates. Nat Commun. 2018;9: 3013. doi.org/10.1038/s41467-018-05419-7

19. Avise JC. Phylogeography: retrospect and prospect. J Biogeog. 2009;36: 3–15. doi.org/10.1111/j.1365-2699.2008.02032.x

20. Brusca RC. A brief geologic history of northwestern Mexico. vers. 12, at http://rickbrusca.com/http:www.rickbrusca.com_index.html/research.html). 2015; pp.1-84.

21. Umhoefer PJ, Dorsey RJ Translation of terranes: lessons from central Baja California, Mexico. Geology. 1997;25: 1007–1010. doi.org/10.1130/0091-7613(1997)025<1007:TOTLFC>2.3.CO;2

22. Calmus T, Pallares C, Bellon H, Benoit M, Maury R, Aguillón-Robles A, Cotton J. Temporal geochemical evolution of Neogene volcanism in northern Baja California (27°-30°N): insights on the origin of post-subduction magnesian andesites. Lithos. 2008;105: 162–180. doi.org/10.1016/j.lithos.2008.03.004

23. Karrig R, Jensky W. The proto Gulf of California. Earth Planet. Sci. Lett. 1972;17: 169–174. doi.org/10.1016/0012-821X(72)90272-5

24. Grismer L. Evolutionary biogeography on Mexico’s Baja California peninsula: A synthesis of molecules and historical geology. Proc Nat Acad Sci. 2000;97: 14017– 14018. doi.org/10.1073/pnas.26050969

25. Riddle BR, Hafner DJ, Alexander LF, Jaeger JR. Cryptic vicariance in the historical assembly of a Baja California peninsular desert biota. Proc Nat Acad Sci. 2000;97: 14438–14443. doi.org/10.1073/pnas.250413397

26. Dolby GA, Bennett SEK, Lira-Noriega A, Wilder BT, Munguía-Vega A. Assessing the geological and climatic forcing of biodiversity and evolution surrounding the Gulf of California. J Southwest. 2015;57: 391–455. doi: 10.1353/jsw.2015.0005

27. Klimova A, Hoffman JI, Gutierrez-Rivera JN, Leon de la Luz J, Ortega-Rubio A. Molecular genetic analysis of two native desert palm genera, *Washingtonia* and *Brah*ea, from the Baja California Peninsula and Guadalupe Island. Ecol Evol. 2017;7: 4919– 4935. doi.org/10.1002/ece3.3036

28. Turner R, Bowers J, Burgess T. Sonoran Desert Plants: An ecological atlas. Tucson: University of Arizona Press; 1995.

29. Riemann H, Ezcurra E. Endemic regions of the vascular flora of the peninsula of Baja California, Mexico. J Veg Sci. 2007;18: 327–336. doi.org/10.1111/j.1654-1103.2007.tb02544.x

30. Wilder BT, Felger RS, Ezcurra E. Controls of plant diversity and composition on a desert archipelago. PeerJ. 2019;7:e7286. doi 10.7717/peerj.7286

31. Riginos C. Cryptic vicariance in Gulf of California fishes parallels vicariant patterns found in Baja California mammals and reptiles. Evolution. 2005;59: 2678–2690. doi.org/10.1111/j.0014-3820.2005.tb00979.x

32. Nason, JD, Hamrick JL, Fleming TH. Historical vicariance and postglacial colonization effects on the evolution of genetic structure in *Lophocereus*, a Sonoran Desert columnar cactus. Evolution. 2002;56: 2214–2226. doi: 10.1111/j.0014-3820.2002.tb00146.x

33. Mulcahy D, Macey R. Vicariance and dispersal form a ring distribution in nightsnakes around the Gulf of California. Mol Phylogenet Evol. 2009;53: 537–546. doi.org/10.1016/j.ympev.2009.05.037

34. Martínez-Noguez JJ, León de la Luz JL, Delgadillo Rodríguez J, García-De León FJ Phylogeography and genetic structure of an iconic tree of the Sonoran Desert, the Cirio (*Fouquieria columnaris*), based on chloroplast DNA, Biol J Linn Soc. 2020;130: 433–446. doi.org/10.1093/biolinnean/blaa065

35. Yetman D, Búrquez, A. Hultine K, Sanderson M. The Saguaro: A natural history. Tucson: University of Arizona Press; 2020.

36. Bustamante E, Casas A, Búrquez A. Geographic variation in reproductive success of *Stenocereus thurberi* (Cactaceae): effects of pollination timing and pollinator guild. Am J Bot. 2010;97: 2020–2030. doi: 10.3732/ajb.1000071

37. Bustamante E, Búrquez A, Scheinvar E, Eguiarte LE. Population genetic structure of a widespread bat-pollinated columnar cactus. PLoS ONE. 2016;11: 1–18. doi.org/10.1371/journal.pone.0152329

38. Bustamante E, Búrquez A. Effects of plant size and weather on the flowering phenology of the organ pipe cactus (*Stenocereus thurberi*). Ann Bot. 2008;102: 1019–1030. doi.org/10.1093/aob/mcn194

39. Sosa VJ, Fleming TH. Why are columnar cacti associated with nurse plants? In: Fleming, TH, Valiente-Banuet A. editors. Columnar cacti and their mutualistic: evolution, ecology, and conservation. Tucson; University of Arizona Press. 2002 pp, 306–323.

40. Shaw J, Lickey EB, Beck JT, Farmer SB, Liu W, Miller MJ, Siripun KC, Winder CT, Schilling EE, Small RL. The tortoise and the hare II: relative utility of 21 noncoding chloroplast DNA sequences for phylogenetic analysis. Am J Bot. 2005;92: 142–166. doi: 10.3732/ajb.92.1.142

41. Kroeger TS, Watkins KP, Friso G, van Wijk KJ, Barkan A.A plant-specific RNA-binding domain revealed through analysis of chloroplast group II intron splicing. Proc Nat Acad Sci. 2009;106: 4537–4542. doi:10.1073/pnas.0812503106

42. Edgar RC. MUSCLE: Multiple sequence alignment with high accuracy and high throughput. Nucleic Acids Res. 2004;32: 1792–1797. doi: 10.1093/nar/gkh340

43. Rozas J, Ferrer-Mata A, Sanchez-DelBarrio JC, Guirao-Rico S, Librado P, Ramos-Onsins SE, Sanchez-Gracia A. DnaSP 6: DNA sequence polymorphism analysis of large data sets. Mol Biol Evol. 2017;34(12): 3299–3302, doi.org/10.1093/molbev/msx248.

44. Nei M. Molecular Evolutionary Genetics. New York: Columbia University Press; 1987.

45. Guindon S, Gascuel O. A simple, fast, and accurate algorithm to estimate large phylogenies by maximum likelihood. Syst Biol. 2003;52: 696–704. doi : 10.1080/10635150390235520

46. Ronquist F, Huelsenbeck JP. MrBayes 3: Bayesian phylogenetic inference under mixed models. Bioinformatics. 2003;19: 1572–1574. doi : 10.1093/bioinformatics/btg180

47. Arias S, Terrazas T, Arreola-Nava HJ, Vázquez-Sánchez M, Cameron KM. Phylogenetic relationships in *Peniocereus* (Cactaceae) inferred from plastid DNA sequence data. J Plant Res. 2005;118: 317–328. doi : 10.1007/s10265-005-0225-3

48. Hernández-Hernández T, Hernández HM, De-Nova JA, Puente R, Eguiarte LE, Magallón S. Phylogenetic relationships and evolution of growth form in Cactaceae (Caryophyllales, Eudicotyledoneae). Am J Bot. 2011;98: 44–61. doi: 10.3732/ajb.1000129.

49. Plume OS, Shannon CK, Tel-Zur N, Cisneros A, Schneider B, Doyle JJ. Testing a hypothesis of intergeneric allopolyploidy in vine cacti (Cactaceae: Hylocereeae). Syst Bot. 2013;38: 737–751. doi.org/10.1600/036364413X670421

50. Bandelt HJ, Forster P, Röhl A. Median-joining networks for inferring intraspecific phylogenies. Mol Biol Evol. 1999;16: 37–48. doi: 10.1093/oxfordjournals.molbev.a026036

51. Cullingham CI, Merrill EH, Pybus MJ, Bollinger TK, Wilson GA, Coltman DW. Broad and fine-scale genetic analysis of white-tailed deer populations: Estimating the relative risk of chronic wasting disease spread. Evol Appl. 2011;4: 116–131. doi: 10.1111/j.1752-4571.2010.00142.x

52. Excoffier L, Lischer HEL. Arlequin suite ver 3.5: A new series of programs to perform population genetics analyses under Linux and Windows. Mol Ecol Resour. 2010;10: 564–567. doi.org/10.1111/j.1755-0998.2010.02847.x

53. Slatkin M. Isolation by distance in equilibrium and non-equilibrium populations. Evolution. 1993;47: 264–279. doi.org/10.1111/j.1558-5646.1993.tb01215.x

54. Manni F, Guerard E, Heyer E. Geographic patterns of (genetic, morphologic, linguistic) variation: How barriers can be detected by using Monmonier’s algorithm. Hum Biol. 2004;76: :173–190. doi: 10.1353/hub.2004.0034. digitalcommons.wayne.edu/humbiol/vol76/iss2/1

55. Pritchard JK, Stephens M, Donnelly P. Inference of population structure using multilocus genotype data. Genetics. 2000;155: 945–959. doi.org/10.1093/genetics/155.2.945

56. Hey J, Nielsen R. Multilocus methods for estimating population sizes, migration rates and divergence time, with applications to the divergence of *Drosophila pseudoobscura* and *D. persimilis*. Genetics. 2004;167: 747–760. doi: 10.1534/genetics.103.024182

57. Hey J. Isolation with migration models for more than two populations. Mol Biol Evol. 2010;27: 905–920. doi.org/10.1093/molbev/msp296

58. Chung Y. Recent advances in Bayesian inference of isolation-with-migration models. Genom Inform. 2019;17(4): e37. doi: 10.5808/GI.2019.17.4.e37

59. Zhao YJ, Gong X. Genetic divergence and phylogeographic history of two closely related species (*Leucomeris decora* and *Nouelia insignis*) across the ‘Tanaka Line’ in Southwest China. BMC Evol Biol. 2015;15:134. doi.org/10.1186/s12862-015-0374-5

60. Ossowski S, Schneeberger K, Lucas-Lledó JI, Warthmann N, Clark RM, Shaw RG, Weigel DLM. The rate and molecular spectrum of spontaneous mutations in *Arabidopsis thaliana*. Science. 2010;327: 1–9. doi: 10.1126/science.1180677

61. Bouckaert R, Heled J, Kühnert D, Vaughan T, Wu CH, Xie D, Suchard MA, Rambaut A, Drummond AJ. BEAST 2: A Software platform for Bayesian evolutionary analysis. PLoS Comput Biol. 2014;10: e1003537. doi:10.1371/journal.pcbi.1003537

62. Suchard MA, Drummond AJ. Bayesian random local clocks, or one rate to rule them all. BMC Biology. 2010;8: 114. doi.org/10.1186/1741-7007-8-114

63. Hernández-Hernández T, Brown JW, Schlumpberger BO, Eguiarte LE, Magallón S. Beyond aridification: multiple explanations for the elevated diversification of cacti in the New World Succulent Biome. New Phytol. 2014;202:1382–1397. doi.org/10.1111/nph.12752

64. Phillips SJ, Dudík M. Modeling of species distributions with MAXENT: new extensions and a comprehensive evaluation. Ecography. 2008;31: 161–175. doi: 10.1111/j.0906-7590.2008.5203.x

65. Fick SE, Hijmans RJ. WorldClim 2: new 1-km spatial resolution climate surfaces for global land areas. Int J Climatol. 2017;37: 4302–4315. doi.org/10.1002/joc.5086

66. Evanno G, Regnaut S, Goudet J. Detecting the number of clusters of individuals using the software STRUCTURE: A simulation study. Mol Ecol. 2005;14: 2611– 2620. doi: 10.1111/j.1365-294X.2005.02553.x

67. Wood D, Fisher R, Reeder T. Novel patterns of historical isolation, dispersal, and secondary contact across Baja California in the Rosy Boa (*Lichanura trivirgata)*. Mol Phylogen Evol. 2008;46(2): 484–502. doi: 10.1016/j.ympev.2007.11.014

68. Rebman JP, Roberts NC. Baja California Plant Field Guide. 3rd edition. San Diego: San Diego Natural History Museum; 2012.

69. Ramirez-Acosta J, Castellanos A, Arnaud G, Breceda A, Rojas-Soto O. Conservation of endemic terrestrial vertebrates in the protected areas of the Baja California peninsula, Mexico. Nat. Areas J. 2012;32(1): 15–30.

70. Bernardi G, Findley L, Rocha-Olivares A. Vicariance and dispersal across Baja California in disjunct marine fish populations. Evolution 2003;57: 1599–1609. doi.org/10.1111/j.0014-3820.2003.tb00367.x

71. Breslin PB, Wojciechowski MF, Majure LC. Molecular phylogeny of the Mammilloid clade (Cactaceae) resolves the monophyly of *Mammillaria*. Taxon. 2021;70: 308–323. doi.org/10.1002/tax.12451

72. Martínez-Noguez JJ, León de la Luz JL, Delgadillo Rodríguez J, García-De León FJ, Phylogeography and genetic structure of an iconic tree of the Sonoran Desert, the Cirio (Fouquieria columnaris), based on chloroplast DNA. Biol J Linn Soc. 2020;130(3):433–446 doi.org/10.1093/biolinnean/blaa065

73. Croizat L, Nelson G, Rosen D. Centers of origin and related concepts. Syst Zool. 1974;23: 265–287.

74. Kropf M, Comes H, Kadereit J. Long-distance dispersal vs. vicariance: The origin and genetic diversity of alpine plants in the Spanish Sierra Nevada. New Phytol. 2006;172: 169–184. doi: 10.1111/j.1469-8137.2006.01795.x

75. Pérez M, Téo M, Zappi D, Taylor N, Morales E. Isolation, characterization, and cross-species amplification of polymorphic microsatellite markers for *Pilosocereus machrisii* (Cactaceae). Am J Bot. 2011;98(8): 204–206. doi: 10.3732/ajb.1100033

76. Cody M, Moran R, Thompson H. 1983. The plants. *In*: Case TJ, Cody ML, editors. Island Biogeography in the Sea of Cortez. Berkeley: University of California Press; 49–97.

77. Brousseau L, Fine PV, Dreyer E, Vendramin G, Scotti I. Genomic and phenotypic divergence unveil microgeographic adaptation in the Amazonian hyperdominant tree *Eperua falcata* Aubl. (Fabaceae). Mol Ecol. 2021;30: 1136–1154. doi.org/10.1111/mec.15595

78. Cruzado-Vargas AL, Blanco-García A, Lindig-Cisneros R, Gómez-Romero M, Lopez-Toledo L, de la Barrera E, Sáenz-Romero C. Reciprocal common garden altitudinal transplants reveal potential negativei of climate change on *Abies religiosa* populations in the Monarch Butterfly Biosphere Reserve overwintering sites. Forests. 2021;12: 69. doi.org/10.3390/f12010069

79. Sanderson MJ, Búrquez A, Copetti D, McMahon MM, Zeng Y, Wojciechowski MF Origin and diversification of the saguaro cactus (*Carnegiea gigantea*): a within-species phylogenomic analysis, Syst. Biol. 2022; 71: 1178–1194. doi.org/10.1093/sysbio/syac017

80. de Villemereuil P, Gaggiotti O, Mouterde M, Till-Bottraud I. Common Garden experiments in the genomic era: new perspectives and opportunities. Heredity. 2016;116: 249–254. doi.org/10.1038/hdy.2015.93

81. Búrquez A, Martínez-Yrizar A, Felger RS, Yetman D. Vegetation and habitat diversity at the southern edge of the Sonoran Desert. In: Robichaux RH, editor. Ecology of Sonoran Desert plants and plants communities. Tucson: University of Arizona Press; 1999. pp. 36– 67.

82. Felger RS, Johnson MB, Wilson MF. The Trees of Sonora, Mexico. Oxford: Oxford University Press; 2001.

83. Petit RJ, Hampe A. Some evolutionary consequences of being a tree. Ann Rev Ecol Evol Syst. 2006;37: 187–214. doi.org/10.1146/annurev.ecolsys.37.091305.110215

84. de la Luz L LJ, Navarro P, Juan J, Breceda A. transitional xerophytic tropical plant community of the Cape Region, Baja California. J Veget Sci. 2000;11: 555–564. doi.org/10.2307/3246585

85. Taylor NP. *Stenocereus thurberi subsp. littoralis* (K.Brandegee) N. P.Taylor, Cactaceae Consensus Initiatives. 1998 5:13.

86. Gibson AC. The systematics and evolution of subtribe Stenocereinae. 8. Organ pipe cactus and its closest relatives. Cact. Succ. J. (Los Angeles). 1990;62: 13–24.

87. Sexton JP, Hangartner SB, Hoffmann AA. Genetic isolation by environment or distance: which pattern of gene flow is most common? Evolution. 2014;68: 1–15. doi: 10.1111/evo.12258.

88. Godínez-Alvarez H, Valiente-Banuet A. Fruit-feeding behavior of the bats *Leptonycteris curasoae* and *Choeronycteris mexicana* in flight cage experiments: consequences for dispersal of columnar cactus seeds. Biotropica. 2000; 32:552–556. doi.org/10.1111/j.1744-7429.2000.tb00502.x

89. Valiente-Banuet A, Arizmendi MC, Rojas-Martínez A, Domínguez-Canseco L. Ecological relationships between columnar cacti and nectar-feeding bats in Mexico. J Trop Ecol. 1996;12: 103–119. doi:10.1017/S0266467400009330

90. Medellín RA, Rivero M, Ibarra A, de la Torre JA, González-Terrazas TP, Torres-Knoop L, Tschapka M. Foraging distances of *Leptonycteris yerbabuenae* (Chiroptera: Phyllostomidae) in Sonora determined by fluorescent dust. J Mammal. 2018;99(3): 306–311. doi.org/10.1093/jmammal/gyy016

91. Wolf BO, Martinez Del Rio C. Use of saguaro fruit by white-winged doves: isotopic evidence of a tight ecological association. Oecologia. 2000;124: 536–543. doi: 10.1007/s004420000406.

92. Hewitt G. The genetic legacy of the Quaternary ice ages. Nature. 2000;405: 907– 913. doi.org/10.1038/35016000

93. Ossa CG, Montenegro P, Larridon I, Pérez F. Response of xerophytic plants to glacial cycles in southern South America. Ann Bot. 2019;124: 15–26. doi.org/10.1093/aob/mcy235

94. Rojas-Martínez A, Valiente-Banuet A, Arizmendi MC, Alcántara-Eguren A, Arita HT. Seasonal distribution of the long-nosed bat (*Leptonycteris curasoae*) in North America: does a generalized migration pattern really exist? J Biogeogr. 1999;26: 1065–1077. doi.org/10.1046/j.1365-2699.1999.00354.x

95. Abedi-Lartey M, Dechmann DKN, Wikelski M, Scharf AK, Fahr J. Long-distance seed dispersal by straw-coloured fruit bats varies by season and landscape. Global Ecology and Conservation. 2016;7: 12–24. doi.org/10.1016/j.gecco.2016.03.005

96. Ramirez J. Population genetic structure of the lesser long-nosed bat (*Leptonycteris yerbabuenae)* in Arizona and Mexico. M. Sc. Diss. University of Arizona. 2011.

97. Frick WF, Heady PA, Earl AD, Arteaga MC, Cortés-Calva P, Medellín RA. Seasonal ecology of a migratory nectar-feeding bat at the edge of its range. J Mammal. 2018;99: 1072–1081. doi: 10.1093/jmammal/gyy088.

98. Arteaga MC, Medellín RA, Luna-Ortíz PA, Heady PA, Frick WF. Genetic diversity distribution among seasonal colonies of a nectar-feeding bat (*Leptonycteris yerbabuenae*) in the Baja California Peninsula. Mammal Biol. 2018;92: 78–85. doi.org/10.1016/j.mambio.2018.04.00.

99. Sahley CT, Horner MA, Fleming TH. Flight speeds and mechanical power outputs of the nectar-feeding bat, *Leptonycteris curasoae* (Phyllostomidae: Glossophaginae). J Mammal. 1993;74(3): 594–600. doi.org/10.2307/1382278

100. Buecher DC, Sidner R. Long distance commutes by lesser long-nosed bats (Leptonycteris yerbabuenae) to visit residential hummingbird feeders. In: Gottfried, GJ, Folliott PF, Gebow BS, Eskew LG, Collins LC, editors. Biodiversity and management of the Madrean Archipelago III. Proceedings. RMRS-P-67. Fort Collins: U.S. Department of Agriculture, Forest Service, Rocky Mountain Research Station; 2013. pp. 427–433.

101. Goldshtein A, Handel M, Eitan O, Bonstein A, Shaler T, Collet S, Greif S, Medellín RA, Emek Y, Korman A, Yovel Y. Reinforcement learning enables resource partitioning in foraging bats. Curr Biol. 2020;30: 4096–4102. doi.org/10.1016/j.cub.2020.07.079.

102. Morales-Acuña E, Torres CR, Linero-Cueto JR. Surface wind characteristics over Baja California Peninsula during summer. Reg. Stud. Mar. Sci. 2019;29: 100654. doi.org/10.1016/j.rsma.2019.100654

103. Åkesson S, Bianco G. Wind-assisted sprint migration in northern swifts. iScience. 2021;24: 102474 doi.org/10.1016/j.isci.2021.102474.

104. Dechmann DKN, Wikelski M, Ellis-Soto D, Safi K, Teague O’Mara M. Determinants of spring migration departure decision in a bat. Biol Lett. 2017;13: 20170395. http://dx.doi.org/10.1098/rsbl.2017.0395.

